# Attenuated *Mycobacterium tuberculosis* Vaccine Protection in a Low Dose Murine Challenge Model

**DOI:** 10.1101/2023.03.17.533226

**Authors:** Samuel J. Vidal, Daniel Sellers, Jingyou Yu, Shoko Wakabayashi, Jaimie Sixsmith, Malika Aid, Julia Barrett, Sage F. Stevens, Xiaowen Liu, Wenjun Li, Courtney R. Plumlee, Kevin B. Urdahl, Amanda J. Martinot, Dan H. Barouch

## Abstract

Bacillus Calmette–Guérin (BCG) remains the only clinically approved tuberculosis (TB) vaccine despite limited efficacy. Preclinical studies of next generation TB vaccines have typically used a murine aerosol challenge model that employs a high supraphysiologic challenge dose. Here we show that the protective efficacy of a live attenuated *Mycobacterium tuberculosis* (Mtb) vaccine candidate ΔLprG markedly exceeds that of BCG in a low dose murine aerosol challenge model. BCG vaccination reduced bacterial loads but failed to prevent establishment or dissemination of infection in this model. In contrast, ΔLprG prevented detectable infection in 61% of mice and resulted in striking anatomic containment of 100% breakthrough infections to a single lung. Protection was partially abrogated in a repeated low dose challenge model, which revealed serum IL-17A, IL-6, CXCL2, CCL2, IFN-γ, and CXCL1 as correlates of protection. These data demonstrate that ΔLprG provides strikingly increased protection compared to BCG, including reduced detectable infection and anatomic containment, in a low dose murine challenge model.

## INTRODUCTION

TB is a leading cause of mortality from infectious disease worldwide with more than 1.5 million deaths in 2020^1^. BCG demonstrates high efficacy against disseminated infection in children, but lifetime efficacy in adults ranges widely between 0% and 80%^2^. Despite this limited efficacy, BCG has remained the sole clinically approved vaccine for nearly a century^3^. Accordingly, the development of next-generation vaccines with favorable safety and improved efficacy profiles relative to BCG is an urgent global health priority^4^. We recently developed a live-attenuated vaccine with a deletion in the *rv1411c-rv1410c* operon in the virulent Mtb H37Rv strain termed ΔLprG that was well-tolerated in immunocompromised mice and demonstrated greater reductions in bacterial burdens than BCG^5,6^.

The murine challenge model has proved valuable for studying TB pathophysiology and vaccine development. For example, the essential contributions of CD4 T cells^7^ interferon gamma (IFN-γ)^8,9^, and tumor necrosis factor alpha (TNF-α)^10^ were described in mice. Furthermore, the antigens comprising the M72/AS01_E_ TB vaccine candidate were characterized in mice^11,12^.

However, Mtb is likely transmitted by small respiratory droplets containing few bacilli^13,14^, and the widely employed 100 colony forming unit (CFU) murine aerosol challenge model generally fails to recapitulate key hallmarks of human disease including granulomatous inflammation^15^ and heterogeneous infections^16^. Moreover, the protective efficacy of vaccines is limited to moderate reductions in bacterial burdens, complicating the preclinical interpretation of results in this model. Accordingly, the development of preclinical models that are both experimentally tractable and more reminiscent of human disease is important for next generation vaccine development^17^.

Recently, a lower dose 1-3 CFU murine aerosol model showed heterogeneous bacterial loads as well as a subset of mice with unilateral infection, and barcoding experiments showed that most cases bilateral infection were driven by dissemination of a single infecting bacterium to the contralateral lung^18^. BCG immunization increased the proportion of mice with unilateral infection and reduced the proportion of mice with detectable infection, although this study required several hundred mice due to the low protective efficacy of BCG^19^. In this study, we evaluated the protective efficacy of BCG and a more potent live attenuated ΔLprG vaccine candidate in the low dose model^5,6^. Compared to BCG, ΔLprG yielded protection from detectable infection in a subset of mice and striking anatomic containment in all animals with breakthrough infection.

## RESULTS

### ΔLprG is immunogenic and protective after 100 CFU challenge

We first sought to confirm the immunogenicity and protective efficacy of the live attenuated ΔLprG vaccine strain against a 100 CFU challenge. We focused these studies in C3HeB/FeJ mice, which exhibit susceptibility to Mtb infection and show granulomas with central caseous necrosis that are reminiscent of human disease^20^. Mice were vaccinated at week 0 with BCG or ΔLprG, and we measured purified protein derivative (PPD)-specific T cell responses in peripheral blood mononuclear cells (PBMCs) at week 2 by intracellular cytokine staining (ICS). BCG did not elicit significant CD4 IFN-γ-secreting (Fig. 1A), CD4 TNF-α- secreting (Fig. 1B), or CD8 IFN-γ-secreting (Fig. 1C) responses in PBMCs at this time point. In contrast, ΔLprG stimulated significant cytokine-secreting T cells responses of all three phenotypes (Fig. 1A-C). Next, we performed multiplexed serum cytokine analysis in naïve and vaccinated mice at week 2.5. Compared to naïve mice, BCG-vaccinated mice showed upregulation of serum cytokines including IFN-γ, TNF-α, and interleukin-17A (IL-17A), among others (Fig. S1A and S1B). ΔLprG also stimulated upregulation of serum cytokines including IFN-γ, TNF-α, and IL-17A, among others (Fig. S1A and S1B). There were no cytokines differentially detected between BCG and ΔLprG, although ΔLprG showed a trend toward greater IL-17A levels (Fig. S1C) as previously reported^5^.

**Fig. 1.**
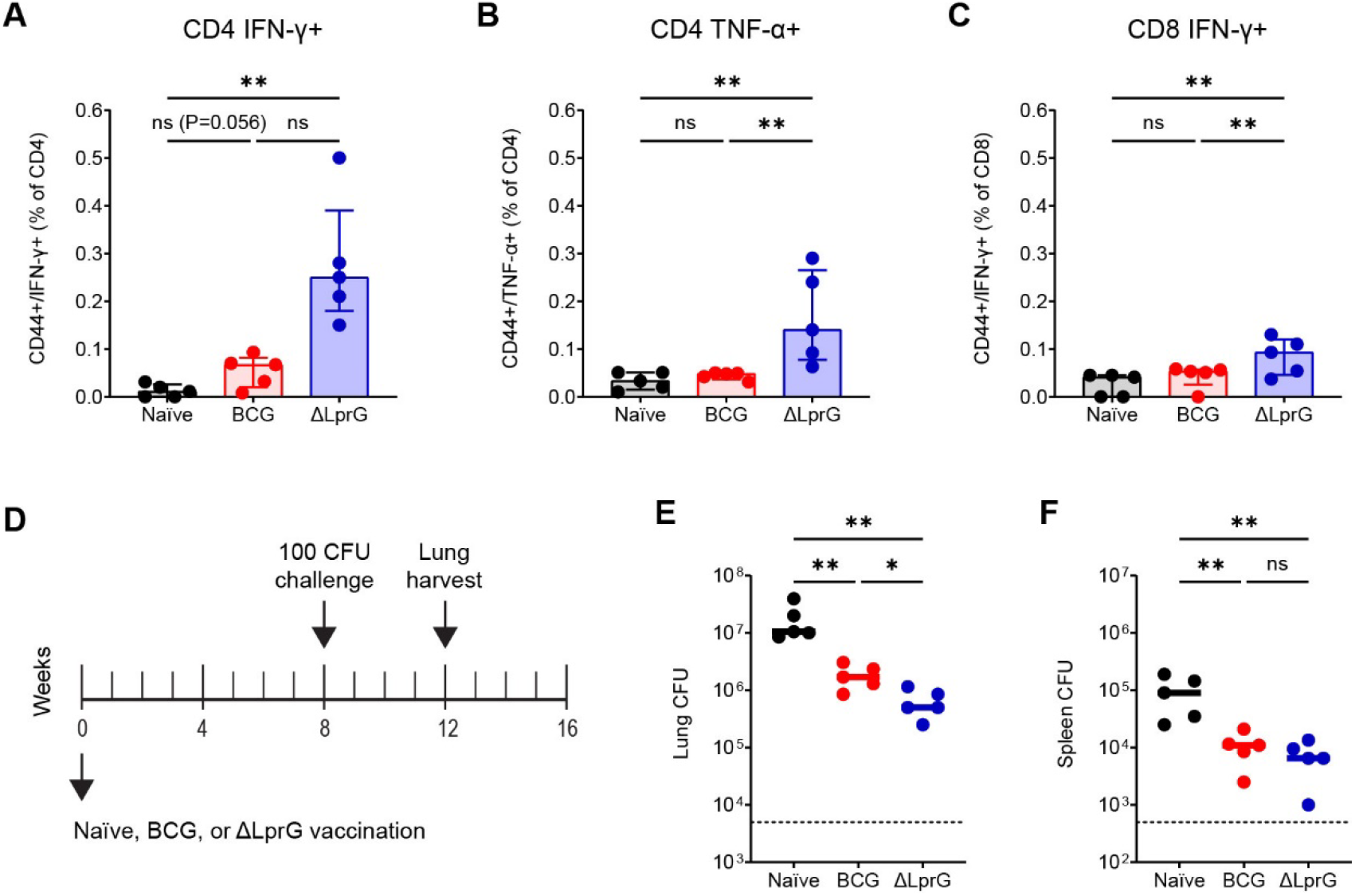
ΔLprG is more immunogenic and protective than BCG following 100 CFU H37Rv challenge in C3HeB/FeJ mice. Groups of C3HeB/FeJ mice were immunized with BCG (n=5) or ΔLprG (n=5) at week 0 followed by PBMC ICS at week 2 to quantify subsets including CD4 IFN-γ+ T cells (A), CD4 TNF-α+ T cells (B), and CD8 IFN-γ+ T cells (C). Challenge study design (D). Groups of C3HeB/FeJ mice were immunized with BCG (n=5) or ΔLprG (n=5) at week 0 followed by 100 CFU H37Rv aerosol challenge at week 8 and lung and spleen harvesting at week 12 for bacterial load quantification. Lung CFU from the challenge study (E). Bottom dotted represents assay LOD of 5,000 CFU. Spleen CFU from the challenge study (F). Bottom dotted line represents assay LOD of 500 CFU. For all panels P values represent pair-wise Mann Whitney U tests. For all panels * represents P<0.05 and ** represents P<0.01.

We next assessed the protective efficacy of BCG and ΔLprG against a 100 CFU H37Rv aerosol challenge. Groups of C3HeB/FeJ mice (n=5 per group) were vaccinated at week 0 with BCG or ΔLprG, challenged at week 8 with 100 CFU of H37Rv by the aerosol route, and lungs and spleens were harvested at week 12 for bacterial load quantification. ΔLprG yielded a greater reduction in bacterial loads in the lung relative to BCG, as we previously reported^5^, and both vaccines reduced bacterial loads in the spleen (Fig. 1E and 1F). Thus, ΔLprG showed a more favorable immunogenicity and protective efficacy profile against 100 CFU challenge.

### Characterization of 1 MID50 challenge in C3HeB/FeJ mice

To adapt the low dose challenge model, we first performed a log_10_ *in vivo* titration of a single-cell suspension H37Rv challenge stock to determine a dose that produced a 60-70% infection rate^18^ (data not shown). We next performed a more focused log_2_-scale *in vivo* dose-finding study, harvesting separately dissected right and left lung lobes 4 weeks after challenge of 30 C3HeB/FeJ mice (n=10 mice per group) with the single-cell suspension H37Rv challenge stock. This study yielded infection rates of 10%, 60%, and 80% with a limit of detection (LOD) of 5 CFU per lung lobe (Fig. 2A-D). These studies demonstrated heterogeneous bacterial loads spanning an approximately 4 log_10_ range and showed that subsets of mice demonstrated unilateral infection with Mtb CFU detected solely in the right or left lung lobe (Fig. 2E and 2F).

**Fig. 2.**
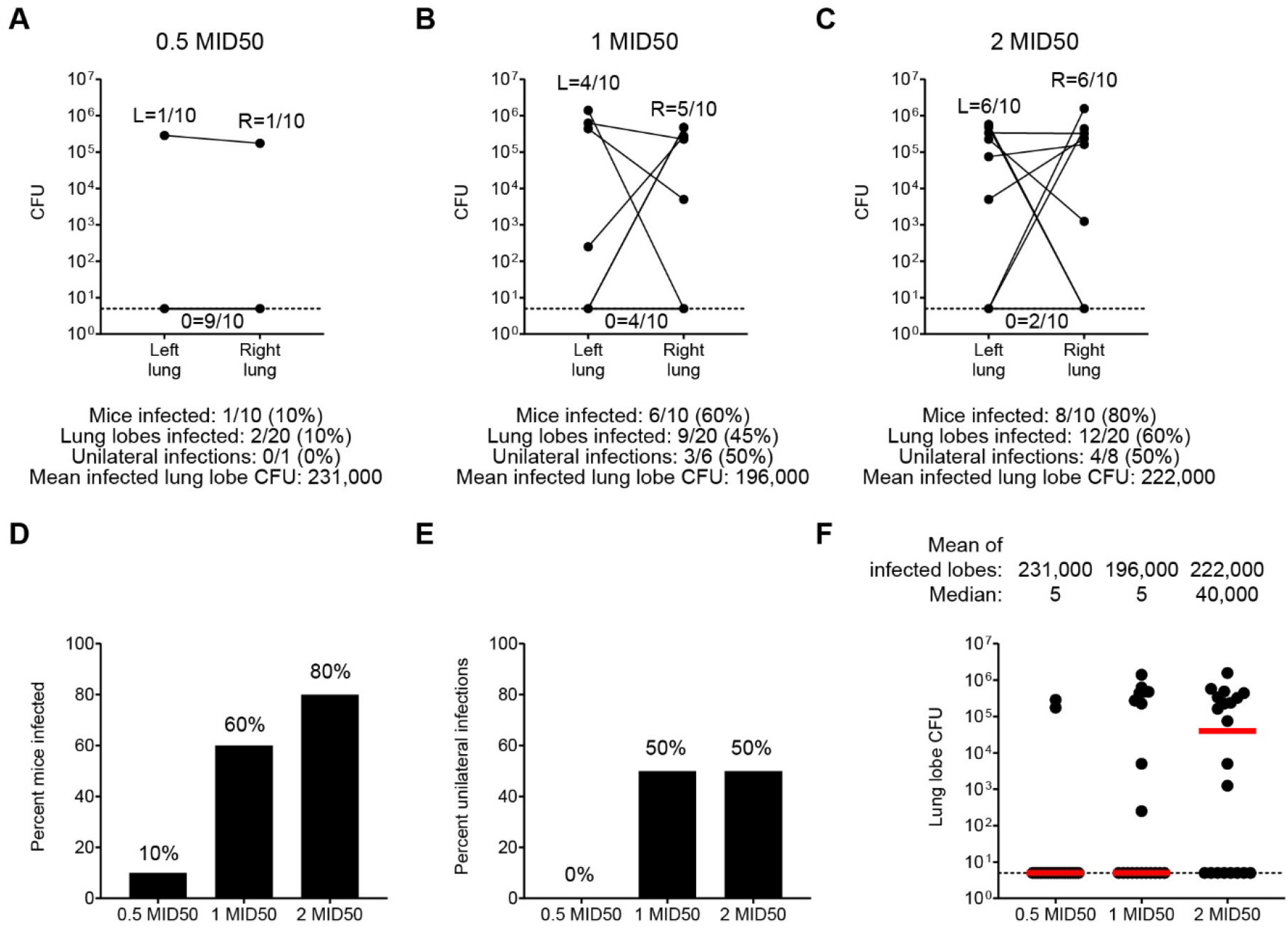
Characterization of 1 MID50 H37Rv challenge in C3HeB/FeJ mice. Single-cell suspension H37Rv challenge stock log_2_-scale *in vivo* dose-finding study (A-C). C3HeB/FeJ mice (n=10 per group) were challenged at week 0 with the indicated challenge stock doses followed by right and left lung lobe harvesting at week 4 for bacterial load quantification. Bottom dotted line represents assay LOD of 5 CFU. Histograms and dot plots summarizing infection rates and bacterial loads shown in A-C (D-F).

In order to formulate a quantitative nomenclature for challenge doses, and given that the intermediate challenge dose yielded a 60% infection rate (Fig. 2B), we termed this challenge dose 1 median infectious dose 50 (1 MID50). We further calculated that the bacterial inoculum required for 100 CFU challenge was approximately 2 log_10_ higher than for the 1 MID50 challenge, and we therefore termed this challenge dose 100 MID50.

Finally, we performed histopathological studies comparing the lung tissues of mice after either 1 MID50 or 100 MID50 H37Rv aerosol challenge. 1 MID50 mice showed fewer granulomas than 100 MID50 mice, and when identified granulomas were smaller and solitary in 1 MID50 compared to 100 MID50 mice (Fig. S2A and S2D). In some cases, 1 MID 50 mice showed only rare foci of alveolitis characterized by increased numbers of alveolar macrophages and minimal expansion of the interstitium with lymphocytes and perivascular lymphocyte cuffing (Fig. S2B and S2E). Acid-fast staining revealed multibacillary (>2 bacilli per macrophage) replication of Mtb in 100 MID50 mice compared to typically paucibacillary macrophage infection in 1 MID50 mice (Fig. S2C and S2F).

### ΔLprG reduces infection and dissemination after 1 MID50 challenge compared to BCG

We next performed vaccine studies with BCG and ΔLprG using 1 MID50 challenge (Fig. 3A). Three cohorts of C3HeB/FeJ mice (n=18 mice per cohort, n=54 total mice) were divided equally into three groups including naïve, BCG, and ΔLprG. Mice were vaccinated at week 0 and underwent a 1 MID50 H37Rv aerosol challenge at week 8. At week 12, right and left lung lobes were dissected separately and bacterial loads were quantified with an LOD of 5 CFU. The pooled infection rate in the naïve group was 13/18 (72%, Fig. 3B), confirming reproducibility of achieving the challenge dose associated with 1 MID50 infection^18^. Among infected naïve animals, there were 4/13 (31%) unilateral infections (Fig. 3B). As before, we observed a broad distribution of bacterial loads with a mean lung lobe bacterial burden of 4.90 log_10_ CFU (Fig. 3B).

**Fig. 3.**
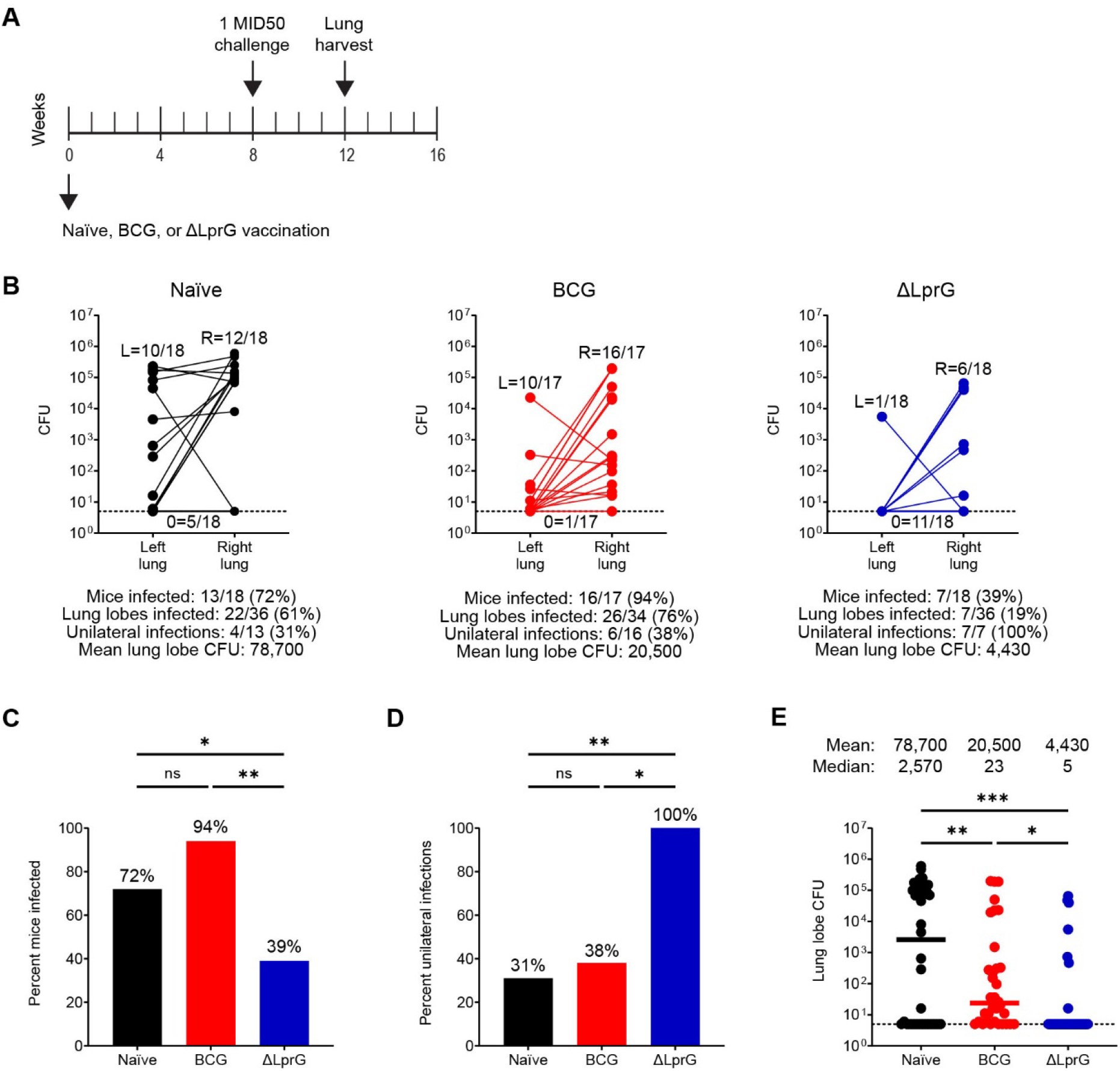
Vaccine protection with BCG and ΔLprG after 1 MID50 H37Rv challenge in C3HeB/FeJ mice. Study design (A). Three cohorts of C3HeB/FeJ mice (n=18 mice per cohort, n=54 mice total) were divided equally into naïve, BCG, and ΔLprG groups. Vaccines were administered at week 0 followed by 1 MID50 H37Rv aerosol challenge at week 8 and right and left lung lobe harvesting at week 12 for bacterial load quantification (B). Bottom dotted line represents an LOD of 5 CFU. Histograms and dot plots summarizing data shown in B (C-E). P values for infection rates between groups represent an exact logistic regression model (C-D). P values for bacterial loads between groups represent a mixed effects negative binomial model (E). For all panels * represents P<0.05, ** represents P<0.01, and *** represents P<0.001.

In the BCG group, we observed an infection rate of 16/17 (95%) including a unilateral infection rate of 6/16 (38%, Fig. 3B). We used an exact logistic regression model to compare both rates of mouse infection (in either one or both lungs) as well as and rates of dissemination (among infected mice to both lungs) between vaccine groups. We observed no difference in the rate of infection or dissemination between the naïve and BCG groups (P=0.12 for both comparisons, exact logistic regression, Fig. 3C and 3D). In addition, we used an ordinal logistic regression model to compare a composite outcome including both infection and dissemination between vaccine groups. We observed no difference in this composite outcome between the naïve and BCG groups (P=0.29, ordinal logistic regression). However, BCG did yield a 0.58 log_10_ reduction in mean lung lobe CFU relative to the naïve group (P<0.001, mixed effects negative binomial, Fig. 3E). Thus, BCG reduced bacterial burdens but failed to abort establishment or dissemination of infection after 1 MID50 challenge.

In contrast to BCG, ΔLprG yielded an infection rate of 7/18 (39%), and all breakthrough mice (7/7, 100%) demonstrated unilateral infection (Fig. 3B). The infection rate in the ΔLprG group was lower compared to both the naïve and BCG groups (P=0.049 and P=0.005, respectively, exact logistic regression, Fig. 3C), and the unilateral infection rate was higher compared to both the naïve and BCG groups (P=0.009 and P=0.014, respectively, exact logistic regression, Fig. 3D). Moreover, the composite outcome of infection and dissemination was lower in the ΔLprG group compared to both the naïve and BCG groups (P=0.002 and P=0.001, respectively, ordinal logistic regression). Finally, the ΔLprG group showed a 1.3 log_10_ reduction in mean lung lobe CFU relative to the naïve group, which also represented a 0.67 log_10_ reduction relative to the BCG group (P<0.001 and P=0.023, respectively, mixed effects negative binomial, Fig. 3E). These data demonstrate that ΔLprG vaccination resulted in a striking reduction in both the establishment and dissemination of infection in this model.

### Repeated 1 MID50 challenge infects most mice and shows greater stringency

In order to better model real-world dynamics including repeated exposure^21,22^, we designed studies incorporating repeated 1 MID50 challenge (Fig. 4A). Two cohorts of C3HeB/FeJ mice (n=18 mice per cohort, n=36 total mice) were divided equally into three groups including naïve, BCG, and ΔLprG. Each cohort was vaccinated week 0, underwent four consecutive 1 MID50 H37Rv challenges at weeks 8, 9, 10, and 11, and at week 15 right and left lungs were dissected separately for bacterial load quantification. For a third cohort (n=18 mice), lungs were fixed in formalin at week 15 for histopathological studies.

**Fig. 4.**
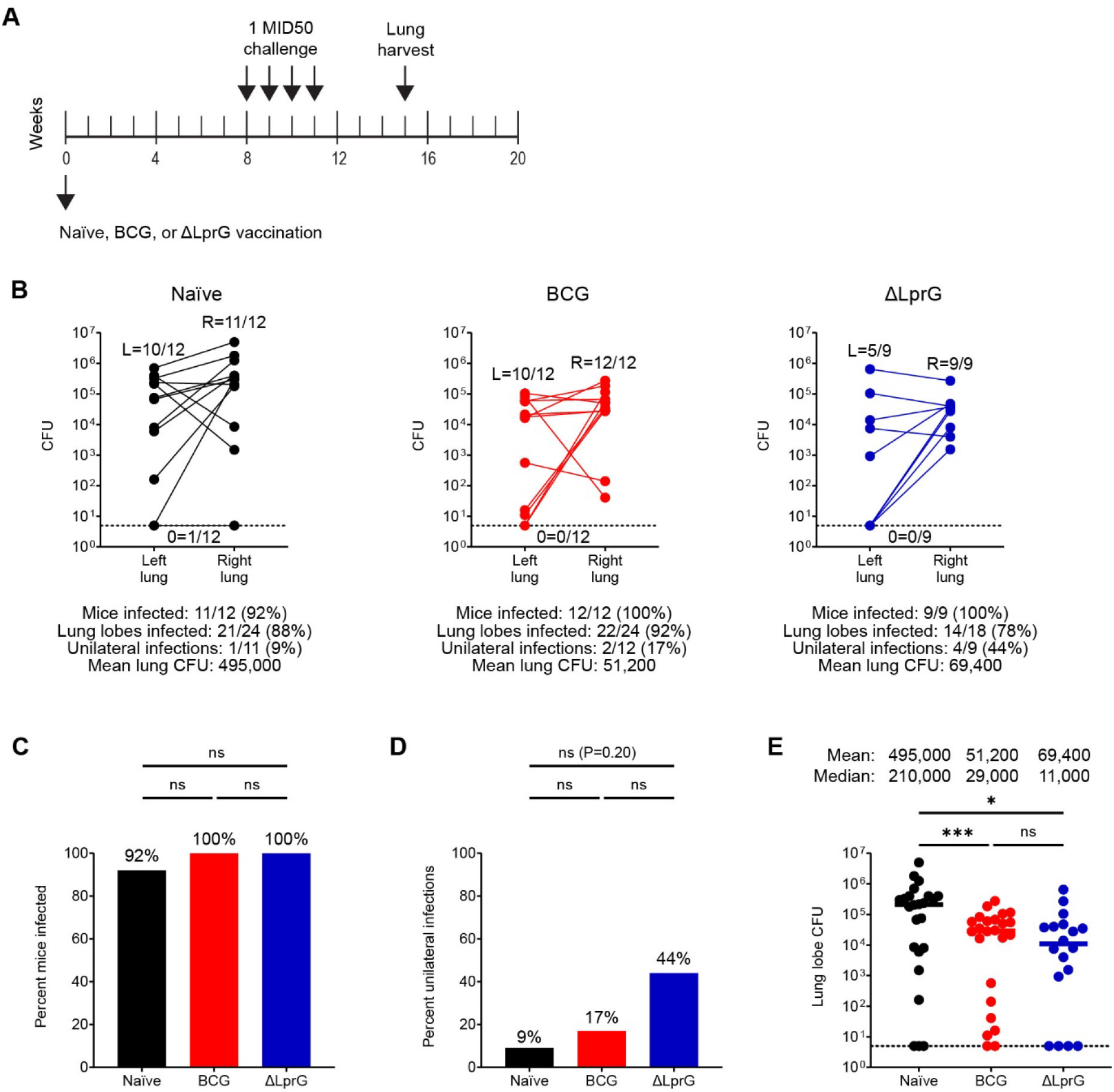
Vaccine protection with BCG and ΔLprG after repeated 1 MID50 H37Rv challenge in C3HeB/FeJ mice. Study design (A). Two cohorts of C3HeB/FeJ mice (n=18 mice per cohort, n=36 mice total) were divided equally into naïve, BCG, and ΔLprG groups. Vaccines were administered at week 0 followed by four weekly 1 MID50 H37Rv aerosol challenges between weeks 8-11 and right and left lung lobe harvesting at week 15 for lung lobe bacterial load quantification (B). Bottom dotted line represents an LOD of 5 CFU. Histograms and dot plots summarizing data shown in B (C-E). P values for infection rates between groups represent an exact logistic regression model (C-D). P values for bacterial loads between groups represent a mixed effects negative binomial model (E). For all panels * represents P<0.05 and *** represents P<0.001.

As expected, we observed an increased proportion of infected mice and a reduced proportion of mice with unilateral infection in the repeated challenge model (Fig. 4B). Specifically, in the naïve group, we observed that 11/12 (92%) of mice became infected, and only 1/11 (9%) mice demonstrated unilateral infection. Similar to single 1 MID50 challenge, there was a wide distribution of bacterial loads. In the naïve group, repeated challenge also yielded increased average lung lobe bacterial loads relative to single low dose challenge (5.69 log_10_ CFU vs. 4.90 log_10_ CFU, respectively, P=0.002, Mann-Whitney U test). Histopathologic studies showed that repeated 1 MID50 challenge granulomas (up to 7 weeks post-initial challenge) in naïve mice were indistinguishable in size and composition from those observed after 100 MID50 challenge (4 weeks post-challenge), consistent with early establishment of infection and prolonged bacterial replication (Fig. S2 and S3A-C).

In the BCG group, we observed an infection rate of 12/12 (100%) including a unilateral infection rate of 2/12 (17%, Fig. 4B). Thus, BCG vaccination did not reduce the rate of infection or increase the rate of unilateral infection (Fig. 4C-D). In contrast, BCG showed a substantial 0.99 log_10_ reduction in lung lobe bacterial loads compared to the naïve group (Fig. 4B and 4E, P<0.001, mixed effects negative binomial). In the ΔLprG group, we observed an infection rate of 9/9 (100%) including a unilateral infection rate of 4/19 (44%, Fig. 4B). In contrast to single 1 MID50 challenge, following repeated challenge ΔLprG did not reduce infection rates relative to the naïve and BCG groups (Fig. 4C). However, we observed a trend towards an increased proportion of mice with unilateral infection relative to the naïve (44% vs. 11%) and BCG (44% vs. 17%) groups (Fig. 4D). Finally, the ΔLprG group showed a 0.85 log_10_ reduction in average lung lobe bacterial lobes relative to the naïve group (Fig. 4B and 4E, P=0.016, mixed effects negative binomial model). Mice vaccinated with both BCG and ΔLprG had fewer and smaller granulomas than naïve mice after repeated low dose challenge, consistent with a delay in acquisition of disease or greater control of bacterial replication (Fig. S3). Increased numbers of lymphocytes as well as increased perivascular lymphocytic cuffing were observed in granulomas from vaccinated mice.

### Repeated 1 MID50 challenge facilitates correlates of protection analyses

We evaluated post-vaccination, pre-challenge serum cytokine levels among one cohort of repeated low dose challenge mice in which all (6/6, 100%) naïve animals became infected and compared them with post-challenge whole-lung CFU on a per-mouse basis. We observed that 7 of 35 assayed post-vaccination serum cytokines were negatively correlated with post-challenge bacterial loads (Fig. 5). Notably, the serum cytokine most correlated with reductions in bacterial load was IL-17A, a molecule that we and others have found to be correlated with protection in both mouse^5^ and macaque^23^ vaccine studies as well as natural infection studies in macaques^24^ and humans^25^. Interestingly, 4 of the remaining 6 correlates of protection were macrophage-secreted cytokines. These included IL-6 and CXCL2 as we previously described in the 100 MID50 challenge model^5^ as well as others including CCL2 and CXCL1 which to our knowledge have not previously been described as correlates of TB vaccine protection.

**Fig. 5.**
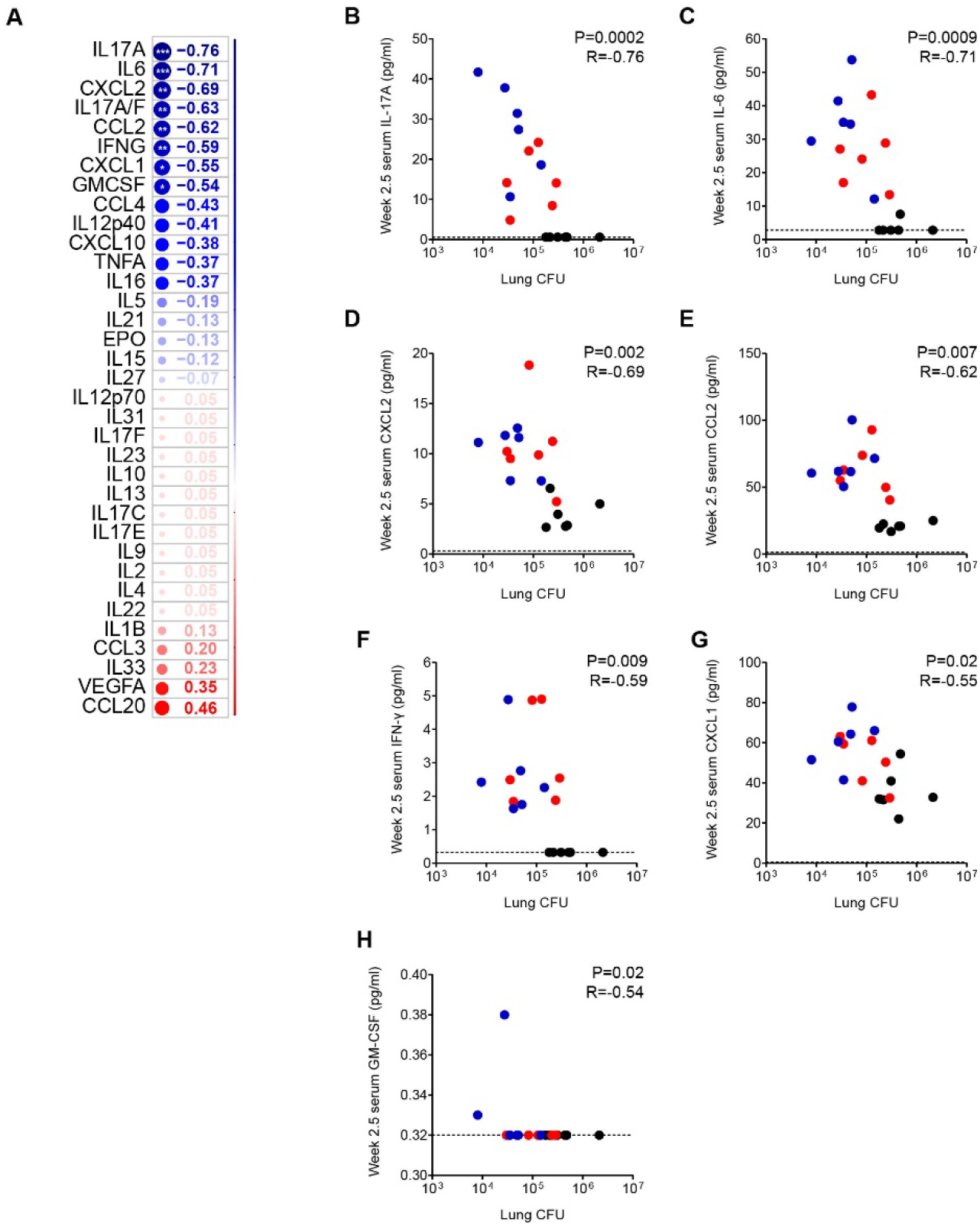
Pre-challenge serum cytokine levels correlated with bacterial loads among naïve and BCG- and ΔLprG-vaccinated C3HeB/FeJ mice after four weekly 1 MID50 H37Rv challenges. A cohort of C3HeB/FeJ mice (n=18) was divided into naïve (n=6), BCG (n=6), and ΔLprG (n=6) groups. Vaccines were administered at week 0 and serum cytokines were measured at week 2.5. The cohort then underwent four weekly 1 MID50 H37Rv aerosol challenges between weeks 8-11 and right and left lung lobes harvested at week 15 for lung lobe bacterial load quantification. Two-tailed Spearman correlation coefficients comparing the serum expression levels of 35 cytokines measured at week 2.5 with whole-mouse bacterial loads measured at week 15 (A). * represents P<0.05, ** represents P<0.01, and *** represents P<0.001. Individual two-tailed Spearman correlation plots for IL-17 (B), IL-6 (C), CXCL2 (D), CCL2 (E), IFN-γ (F), CXCL1 (G), and GM-CSF (H).

## DISCUSSION

Preclinical studies have primarily relied on a 100 MID50 murine aerosol challenge model that does not simulate typical Mtb transmission dynamics found in humans. This model results in diffuse infections with uniformly high bacterial loads that are only moderately reduced by clinical and preclinical vaccines. In this study, we demonstrate striking protective efficacy of the live attenuated ΔLprG preclinical vaccine against a low dose challenge including anatomic containment to a single lung in mice that are infected. In contrast, BCG did not achieve such protection in this model. This reduced dose challenge model provides a promising platform for the evaluation of next generation TB vaccine candidates including ΔLprG.

Mtb transmission typically occurs after repeated exposures, for example in the setting of infected household contacts^21,22^. Indeed, preclinical non-human primate studies are increasingly performed with repeated low dose Mtb challenge^23^. Similarly, repeated low dose exposure is commonly used in the HIV field to test prophylactic interventions^26,27^. In both of these models, repeated challenge creates unique opportunities for measuring correlates of protection^23,26,27^. We therefore developed a repeated low dose Mtb challenge model that resulted in increased challenge stringency and reduced vaccine efficacy compared with the single low dose Mtb challenge model. A correlates of protection analysis suggested that serum IL-17A was a key vaccine correlate of protection^5,23^ and identified additional novel biomarkers of protection including CCL2 and CXCL1. Similar to IL-6 and CXCL2, both CCL2 and CXCL1 are macrophage-secreted as well as associated with disease progression in humans^28,29^. Thus, the repeated low dose challenge model may contribute to the identification of novel correlates of vaccine protection. In future studies, these and other correlates of protection can be studied mechanistically in the single and repeated low dose challenge models. In this context, the unique features of low dose challenge may enable dissection of immunologic mechanisms that modulate establishment and dissemination of Mtb infection in addition to control of bacterial burden in an experimentally tractable small animal model.

In summary, we demonstrate that the attenuated TB vaccine candidate ΔLprG provides strikingly greater protection than BCG against low dose challenge in mice, including prevention of detectable infection in the majority of animals and anatomic containment for all breakthrough infections. We further developed a repeated low dose challenge model that identified novel biomarkers of vaccine protection. These data support a new framework for the preclinical evaluation of next generation TB vaccine candidates that are more protective than BCG as well as studies dissecting the immunologic mechanisms that modulate Mtb infection and dissemination in addition to control of bacterial burden.

## Author Contributions

S.J.V., C.R.P., K.B.U., and D.H.B. conceptualized the study. S.J.V., D.S., J.Y., S.W., J.S., J.B., S.F.S., X.L. performed in the mouse studies and immunology assays. M.A. led the bioinformatics. W.L. performed the statistics. A.J.M. performed the histopathology. S.J.V. and D.H.B. wrote the manuscript. All co-authors reviewed and edited the manuscript.

## Acknowledgements

The authors acknowledge NIH IMPAc-TB contract BAA-NIAID-NIHAI201700104 (K.B.U., D.H.B.) as well as the Bill & Melinda Gates Foundation (INV-050234), the Ragon Institute of MGH, MIT, and Harvard, the Massachusetts Consortium for Pathogen Readiness, and the Musk Foundation (D.H.B.).

## Competing interests

A.J.M. and D.H.B. are co-inventors on a provisional vaccine patent PCT/US2020/059152 (Mycobacterial compositions and biomarkers for use in treatment and monitoring of therapeutic responsiveness). All other authors have no competing interests related to the study.

## Data availability

All data are available in the manuscript or the supplementary material. Correspondence and requests for materials should be addressed to D.H.B.

## METHODS

### Challenge and vaccine strains

H37Rv challenge strain was obtained from the Rubin laboratory (Harvard School of Public Health). BCG vaccine strain was obtained from the Urdahl laboratory (University of Washington). ΔLprG vaccine strain was generated in our laboratory as previously described ^5^. All challenge and vaccine strains were grown in media consisting of Middlebrook 7H9 (BD Difco) containing 10% Middlebrook OADC (BD BBL), 0.5% glycerol (Sigma Aldrich), and 0.05% tween 80 (Sigma Aldrich). For preparation of vaccine stocks, vaccine strains were grown in complete growth medium as above additionally supplemented with 0.05% tyloxapol (Sigma Aldrich). Cells were pelleted twice with resuspension in PBS containing 0.05% tyloxapol and then pelleted a third time with resuspension in PBS containing 0.05% tyloxapol and 15% glycerol. Next cells were passaged though a 40 μm filter and then a 20 μm filter for clump removal, followed by storage at -80°C and titering by agar outgrowth assay. For low dose challenge studies, H37Rv culture was grown to an optical density (OD) of 1 followed by passaging through a 5 μm filter to generate a single-cell suspension, resulting in an approximately 2-log_10_ reduction in titer as measured by agar outgrowth assay.

### Mouse strains, immunizations, and aerosol challenges

Female C3HeB/FeJ mice were obtained from Jackson Laboratory (strain 000658) and stored in sterile conditions at the Harvard School of Public Health. All mouse procedures were performed in accordance with Institutional Animal Care and Use Committee (IACUC) guidelines. All immunizations were performed SC in 8 week-old female mice. 1×10^6^ CFU of tittered vaccine strain was used for all challenge studies. 100 μL of optical density (OD) 1 vaccine strain culture was used for the immunogenicity study. For 50-100 CFU challenge, a Glas-Col instrument was used and challenge stocks were titrated to result in a day 1 lung bacterial load of approximately 50-100 CFU. For low dose challenge, the same instrument was used and singe-cell suspension challenge stocks were titrated to result in a week 4 infection rate of approximately 60-70% in accordance with the Poisson distribution as described previously ^18^.

### PBMC flow cytometry

For immunological studies mice were bled via the submandibular route into RPMI (Gibco) containing 5% EDTA (Invitrogen) route in accordance with IACUC guidance. Buffy layer containing PBMC were isolated via Ficoll (GE Healthcare) centrifugation and transferred into RPMI supplemented with 10% FBS (Gibco) and 1% penicillin-streptomycin (Fisher Scientific). Cells were stimulated with purified protein derivative (PPD, Cedarlane) at 400 ng of peptide per test or media control for 6 hours at 37°C and then rested overnight 4°C. PBMCs were then stained with live/dead and cell surface markers in MACS solution (Miltenyi) supplemented with 2% BSA (Miltenyi) prior to permeabilization with Cytofix/Cytoperm (BD Biosciences) and staining with intracellular markers in Perm/Wash (BD Biosciences). PBMCs were then fixed in 2% formaldehyde and stored at 4°C until flow cytometry on an LSR II flow cytometer (BD Biosciences). Cell surface markers included CD3 (clone 17A2, BD Biosciences), CD19 (clone 6D5, BioLegend), CD4 (clone RM4-5, BioLegend), CD8a (clone 53-6.7, BD Biosciences), CD44 (clone IM7, BD Biosciences), and CD62L (clone MEL-14, BioLegend). Intracellular markers included IFN-γ (clone XMG1.2, BioLegend), IL-2 (clone JES6-5H4, BioLegend), TNF- α (clone MP6-XT22, BioLegend) , IL17-A (clone TC11-18H10.1, BioLegend), and IL-4 (clone 11B11, BD Biosciences).

### Serum cytokine analysis

Plasma levels of 35 biomarkers were tested using U-PLEX Biomarker Group 1 (ms) 35-Plex kits from Meso Scale Discovery (MSD, Rockville, MD). The 35 biomarkers and their detection limits (LLODs) are EPO (4.5pg/mL), GM-CSF (0.16 pg/mL), IFN-γ (0.16 pg/mL), IL-12p70 (48 pg/mL), IL-1β (3.1 pg/mL), IL-2 (1.1 pg/mL), IL-5 (0.63 pg/mL), IL-6 (4.8 pg/mL), KC/GRO (4.8pg/mL), TNF-α (1.3 pg/mL), IL-10 (3.8 pg/mL), IL-13 (2.7 pg/mL), IL-15 (24 pg/mL), IL-17F (24 pg/mL), IL-23 (4.9 pg/mL), IL-27p28/IL-30 (8.7 pg/mL), IL-31 (45 pg/mL), IL-33 (2.2 pg/mL), IL-4 (0.56 pg/mL), VEGF-A (0.77 pg/mL), IL-12/IL-23p40 (1.4 pg/mL), IL-16 (3.6 pg/mL), IL-17A (0.30 pg/mL), IL-17C (2.3 pg/mL), IL-17E/IL-25 (1.6 pg/mL), IL-21 (6.5 pg/mL), IL-22 (1.2 pg/mL), IL-17A/F (0.61 pg/mL), IL-9 (1.4 pg/mL), IP-10 (0.51 pg/mL), MCP-1 (1.4 pg/mL), MIP-1α (0.21 pg/mL), MIP-1β (13 pg/mL), MIP-2 (0.30 pg/mL), MIP-3α (0.10 pg/mL). All above assays were done by Metabolism and Mitochondrial Research Core (Beth Israel Deaconess Medical Center, Boston, MA) following manufacturer’s instruction. The assay plates were read by MESO QUICKPLEX SQ 120 instrument and data were analyzed by Discovery workbench 4.0 software.

### Low dose challenge stock preparation and *in vivo* titration studies

H37Rv challenge strain was obtained and propagated as detailed above. Low dose challenge stock was generated as previously described ^18^. Briefly, H37Rv was grown to an OD of 0.7-0.8. Cultures were passed through a 5 μm filter in order to generate a single cell suspension prior to aliquoting, freezing at -80°C, and tittering. For *in vivo* titration, challenges were performed as detailed above. We first estimated a 100-fold reduction in challenge dose relative to 50-100 CFU challenge and challenged mice with three log_10_ dilutions centered around the predicted low dose challenge dose. This identified a challenge dose with a 44% infection rate and was followed by a secondary dose titration study with three log_2_ dilutions centered around the predicted low dose challenge dose.

### Lung processing and CFU quantification

Mice were euthanized 4 weeks following single 50-100 CFU challenge, 4 weeks following single low dose challenge, or 4 weeks following the final weekly low dose challenge in the repeated low dose challenge model. For 50-100 CFU challenge, both lung lobes were dissected en bloc whereas for low dose challenge right and left lung lobes were dissected separately. Tissues were placed into gentleMACS(tm) M Tubes (Miltenyi Biotec 130-096-335) containing 5 ml of PBS and mechanically dissociated using a gentleMACS(tm) Dissociator (Miltenyi Biotec). Lysates were then plated in serial log_10_ dilutions onto 100×15mm Middlebrook 7H10 plates (Hardy Diagnosatics). In order to achieve an LOD of 5 CFU in low dose challenge studies, we additionally plated 1 ml of lysate onto to two 150×15mm plates containing Middlebrook 7H10 agar (BD Difco), 10% Middlebrook OADC (BD BBL), 0.5% glycerol (Sigma Aldrich), and cycloheximide (Sigma Aldrich) at 100 μg/ml. CFU were counted after a 3 week incubation at 37°C.

### Histopathological studies

Lungs were insufflated with 10% neutral buffer formalin for 48 hours then transferred to 70% ethanol and processed routinely into paraffin blocks for hematoxylin and eosin or Ziehl-Neelsen acid-fast staining. Whole slide scanning (20x) was performed using a Midi II Pannoramic scanner (Epredia) and images evaluated by a boarded veterinary pathologist (AJM) using HALO (Indicalabs).

### Statistical analyses

Pairwise tests on immunogenicity and 50-100 CFU challenge data were performed using GraphPad Prism 9.4.0 software. Heatmaps were generated using the R package pheatmap. Cytokines plasma levels were normalized using the Zscore method implemented in the pheatmap package. The correlation of cytokines with lung CFU was performed using the R package corrplot and Spearman’s method. Statistical evaluation was assessed using a t-test distribution implemented in the R cor.test function. Rates of animal level infection (any lobe vs none) were compared using exact logistic regression models. The composite infection outcomes (none vs. one lobe vs. both lobes) were analyzed using ordinal logistic regression models under the assumption of proportional odds. Between-group differences in bacterial loads were analyzed using mixed effects negative binomial regression models while controlling for lobe side/size (right vs. left). Two lobes of the same animal were considered a cluster, and animals were considered independent from each other.

## SUPPLEMENTARY INFORMATION

Figures S1 to S3

## Supplementary figures and legends

**Fig. S1.**
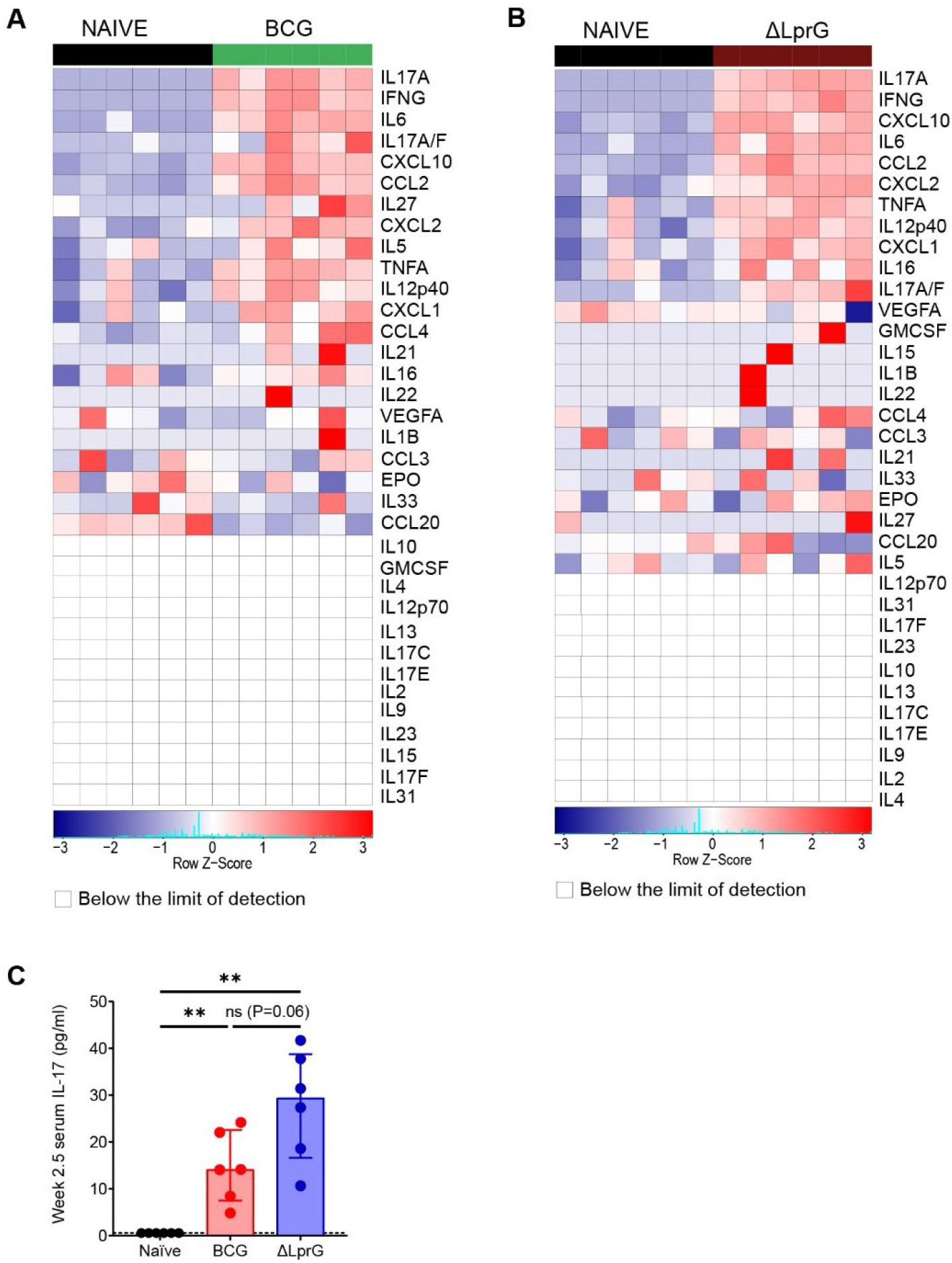
Induction of serum cytokines among naïve and BCG- and ΔLprG-vaccinated C3HeB/FeJ mice. A cohort of C3HeB/FeJ mice (n=18) was divided into naïve (n=6), BCG (n=6), and ΔLprG (n=6) groups. Vaccines were administered at week 0 and serum cytokines were measured at week 2.5. Heat map comparing serum cytokine levels between the naïve and BCG groups (A). Heat map comparing serum cytokine levels between the naïve and ΔLprG groups (B). Serum IL-17A levels among the three groups (C). P values represent pair-wise Mann-Whitney U tests. ** represents P<0.01.

**Fig. S2.**
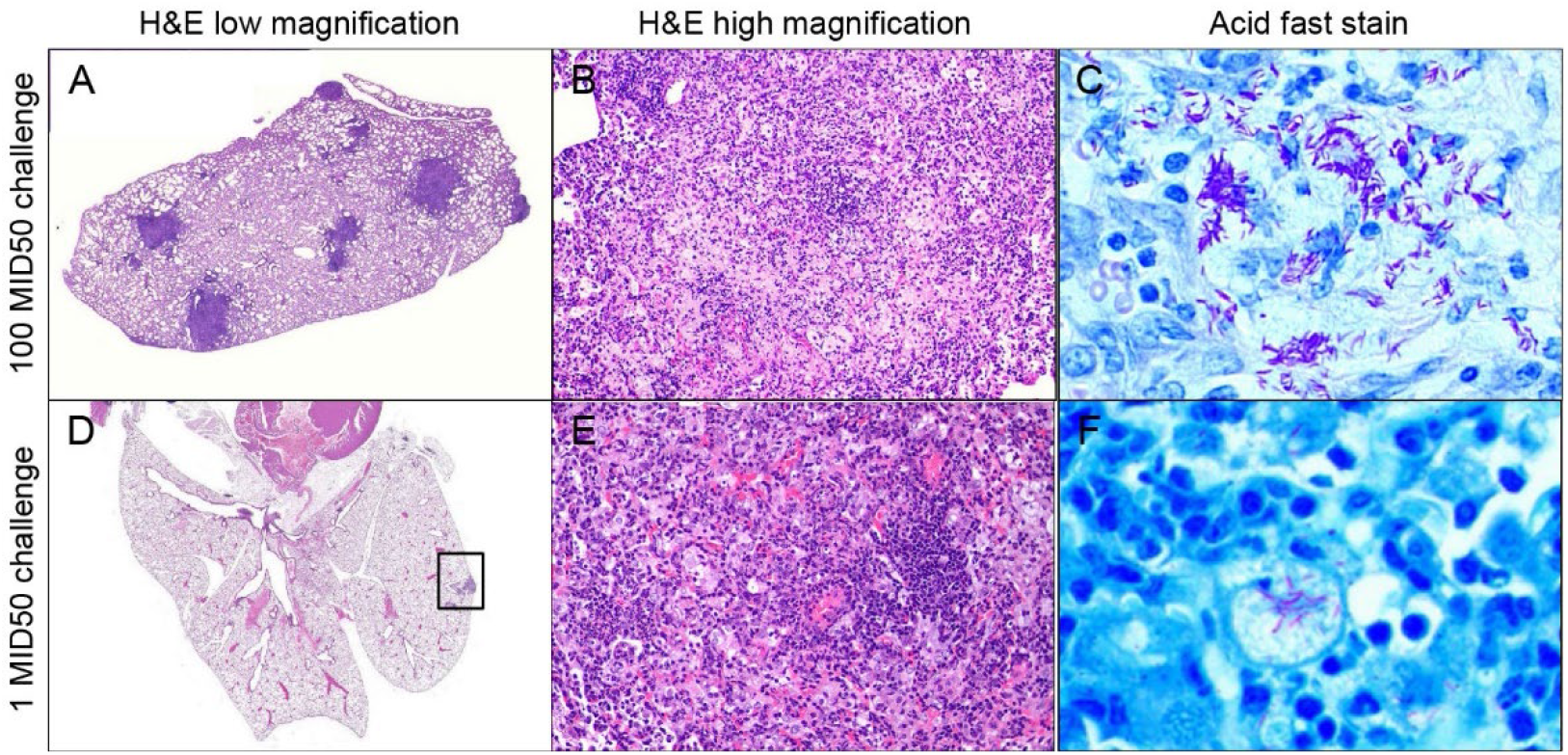
Histopathology of lung tissues from C3HeB/FeJ mice following 1 MID50 or 100 MID50 H37Rv challenge. Representative low-magnification H&E stain (A), high-magnification stain (B), and acid fast stain (C) of lung tissues from C3HeB/FeJ mice 4 weeks after 100 MID50 H37Rv challenge. Representative low-magnification H&E stain (D), high-magnification stain (E), and acid fast stain (F) of lung tissues from C3HeB/FeJ mice 4 weeks after 1 MID50 H37Rv challenge.

**Fig. S3.**
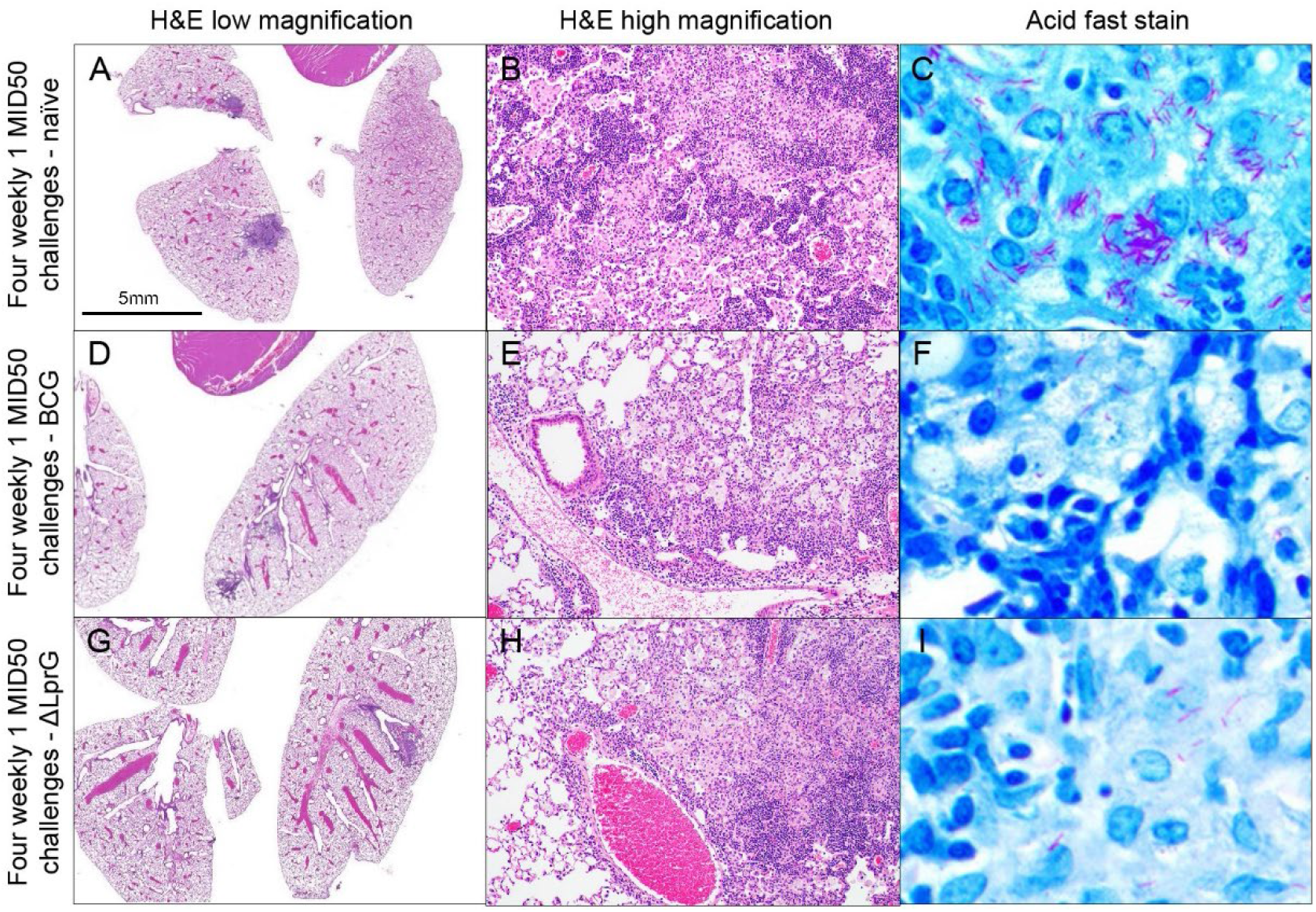
Histopathology of lung tissues from naïve and BCG- and ΔLprG-vaccinated C3HeB/FeJ following four weekly 1 MID50 H37Rv challenges. Representative low-magnification H&E stains (A, D, and G), high-magnification stains (B, E, and H), and acid fast stains (C, F, and I) of lung tissues from C3HeB/FeJ mice 4 weeks after 4 weekly 1 MID50 H37Rv challenges under naïve (A-C), BCG- (D-F), and ΔLprG-vaccinated (G-I) conditions.

